# RNA landscapes of brain tissue and brain tissue-derived extracellular vesicles in simian immunodeficiency virus (SIV) infection and SIV-related central nervous system pathology

**DOI:** 10.1101/2023.04.01.535193

**Authors:** Yiyao Huang, Ahmed Abdelgawad, Andrey Turchinovich, Suzanne Queen, Celina Monteiro Abreu, Xianming Zhu, Mona Batish, Lei Zheng, Kenneth W. Witwer

## Abstract

**Introduction:** Antiretroviral treatment regimens can effectively control HIV replication and some aspects of disease progression. However, molecular events in end-organ diseases such as central nervous system (CNS) disease are not yet fully understood, and routine eradication of latent reservoirs is not yet in reach. Brain tissue-derived extracellular vesicles (bdEVs) act locally in the source tissue and may indicate molecular mechanisms in HIV CNS pathology. Regulatory RNAs from EVs have emerged as important participants in HIV disease pathogenesis. Using brain tissue and bdEVs from the simian immunodeficiency virus (SIV) model of HIV disease, we profiled messenger RNAs (mRNAs), microRNAs (miRNAs), and circular RNAs (circRNAs), seeking to identify possible networks of RNA interaction in SIV infection and neuroinflammation.

**Methods:** Postmortem occipital cortex tissue were collected from pigtailed macaques: uninfected controls and SIV-infected subjects (acute phase and chronic phase with or without CNS pathology). bdEVs were separated and characterized in accordance with international consensus standards. RNAs from bdEVs and source tissue were used for sequencing and qPCR to detect mRNA, miRNA, and circRNA levels

**Results:** Multiple dysregulated bdEV RNAs, including mRNAs, miRNAs, and circRNAs, were identified in acute infection and chronic infection with pathology. Most dysregulated mRNAs in bdEVs reflected dysregulation in their source tissues_._ These mRNAs are disproportionately involved in inflammation and immune responses, especially interferon pathways. For miRNAs, qPCR assays confirmed differential abundance of miR-19a-3p, let-7a-5p, and miR-29a-3p (acute SIV infection), and miR- 146a-5p and miR-449a-5p (chronic with pathology) in bdEVs. In addition, target prediction suggested that several circRNAs that were differentially abundant in source tissue might be responsible for specific differences in small RNA levels in bdEVs during SIV infection.

**Conclusions:** RNA profiling of bdEVs and source tissues reveals potential regulatory networks in SIV infection and SIV-related CNS pathology.

## Introduction

Human immunodeficiency virus (HIV) infection can lead to chronic immune activation and inflammation. Antiretroviral treatment (ART) regimens can effectively control HIV replication, decrease the viral load in the plasma to an undetectable level, and reduce disease progression. However, ART does not eliminate the latent viral reservoir, and end-organ diseases continue to affect people living with HIV (PLWH) (Marsh Sung and M. Margolis, 2018; Siliciano and Greene, 2011). The central nervous system (CNS) can be damaged by chronic inflammation, leading to HIV-associated neurocognitive disorders (HAND) (Smail and Brew, 2018; Tedaldi et al., 2015). Therefore, detecting HIV neuropathology and changes to the brain viral reservoir is of great importance to reveal potential mechanisms of HIV CNS infection and possible HIV cure. SIV-infected models are of great value in the study of HIV viral pathogenesis, eradication, treatment, and vaccine development. Here, we studied archived samples from a pigtailed macaque (*Macaca nemestrina)* model in which subjects are dually inoculated with neurovirulent molecular clone SIV/17E-Fr and an immunosuppressive CD4+ T cell-depleting SIV swarm (B670), recapitulating classic HIV CNS pathology (Beck et al., 2018; MC et al., 1999).

Extracellular vesicles (EVs) are nano-sized membranous vesicles released by most cells. A growing body of evidence suggests broad involvement of EVs in HIV disease pathogenesis. EVs share signaling and biogenesis pathways with HIV, affect virus entry and budding, carry HIV proteins, and transfer molecules with pro-viral and anti-viral effects to recipient cells (Madison and Okeoma, 2015; Stenovec et al., 2019). Much like retroviruses, EVs shuttle bioactive molecules including RNAs and proteins, between cells and are abundant in biofluids (Dreyer and Baur, 2016) like plasma and CSF. However, the most critical site of EV actions may be in the tissue of origin. Tissue-specific EVs can be used to assess the health of the tissue of origin (such as the brain) and may also betray disease mechanisms in the periphery (Li et al., 2020). Recent advances by our group and others allow us to separate and characterize brain tissue-derived EVs (bdEVs) (Huang et al., 2020a) with high rigor. Exploring the molecular components of bdEVs may hint at mechanisms of CNS pathogenesis in the brain (Huang et al., 2022a, 2020b).

Previous studies investigating differences in linear RNAs (mainly messenger RNAs, (mRNAs) (Biswas et al., n.d.; Renga et al., 2012), and microRNAs (miRNAs) (Biswas et al., 2019; Chettimada et al., 2020; Lu et al., 2017; Marques de Menezes et al., 2020; Sun et al., 2012a; Witwer et al., 2011)) in HIV have yielded insights into pathogenic mechanisms that may involve RNA. Circular RNAs (circRNAs) may also contribute towards regulation of HIV replication as nodes in “competing endogenous RNA” (ceRNA) networks (Zhang et al., 2018a). circRNAs are covalently closed single-stranded RNAs that are thought to bind specific miRNAs as “sponges” and thus modulate miRNA posttranscriptional regulatory roles (Liu and Chen, 2022). Since circRNAs are generated from alternative splicing of pre-mRNAs, they also affect linear RNA transcription by competing with canonical RNA splicing from the same pre-mRNAs (Zhang et al., 2020). Like miRNAs, circRNAs are highly stable RNAs and are strong candidates for disease biomarkers. However, regulatory interactions of circRNAs and miRNAs in host immune responses and HIV pathogenesis are largely undetermined, particularly in HIV CNS infection.

We thus assessed potential ceRNA interactions in brain tissue and bdEVs during SIV infection and neuroinflammation using RNA sequencing (RNA-Seq) and quantitative PCR (qPCR) validation. We report associations of mRNA, miRNA, and circRNA in SIV infection and SIV-associated CNS pathology. mRNA differences in brain tissue and bdEVs largely correlated with each other, while miRNA and circRNA findings suggest that several circRNAs in brain tissue may contribute to differential abundance of specific miRNAs in bdEVs. Our study thus proposes novel functions of bdEVs and ceRNAs in HIV infection and HAND.

## Methods

### Tissue sample collection and processing

All samples were from archives of studies approved by the Johns Hopkins University Institutional Animal Care and Use Committee and conducted following the Weatherall Report, the Guide for the Care and Use of Laboratory Animals, and the USDA Animal Welfare Act. Postmortem occipital cortex tissues were obtained from pigtailed macaques (as listed in Table 1) that were not infected (n=6) or dual-inoculated with SIV swarm B670 and clone SIV/17E-Fr (n=16) (Beck et al., 2018; MC et al., 1999). SIV-inoculated groups, including samples collected during acute infection (n=5, 7 days post-inoculation (dpi)) and chronic infection (n=11, 84-101 dpi). According to pathology examination (Mangus et al., 2015; Mankowski et al., 2002), the chronic infection group was divided into subjects without (CP-, n=7) or with CNS pathology (CP+, SIV-encephalitis, n=4). Tissue samples were stored at −80°C.

### Separation of EVs from brain tissue

bdEVs were separated from frozen occipital cortex tissues using our published protocol with minor modifications (Huang et al., 2022a, 2022b, 2020a). Before extraction, a small (∼50_mg) piece of tissue was stored at –80°C for later RNA extraction from brain homogenate (BH). After tissue digestion by 75 U/ml collagenase type 3 (Worthington #CLS-3, S8P18814) for 15 min, the dissociated tissue was spun at 300 × g for 10 min and 2000 × g for 15 min at 4°C. Cell-free supernatant was then filtered through a 0.22 μm filter, and followed by a 10,000 × g spin for 30 min at 4°C. The 10K supernatant was then separated by size exclusion chromatography (SEC) and concentrated.

### Nanoflow cytometry

The concentration and size profile of bdEV preparations were measured by nanoflow cytometry (NFCM; Flow NanoAnalyzer, NanoFCM, Inc.). The instrument was pre-calibrated for concentration and size measurements with 250_nm silica beads and a silica nanosphere cocktail (diameters of 68, 91, 113, and 151_nm), respectively. Both calibration materials were from the manufacturer, NanoFCM. bdEV preparations were diluted as needed (typically 1:200 dilution), and particle events were recorded for 1 minute. Particle numbers and sizes were calculated based on the calibration curve, flow rate, and side scatter intensity.

### Single particle interferometric reflectance imaging (SP-IRIS)

Measurements were performed per manufacturer’s instructions and as previously described (Arab et al., 2021). Briefly, A total of 10 μL bdEVs were diluted in 35 μL incubation buffer (IB). Diluted EVs were then incubated overnight at room temperature on ExoView chips (NanoView Biosciences) printed with anti-human CD81, CD63, CD9, and isotype controls. After incubation, chips were washed with IB 4 times for 3 min and then imaged in the ExoView scanner (NanoView Biosciences, Brighton, MA) by interferometric reflectance imaging detection. Data were analyzed using NanoViewer 2.8.10 Software (NanoView Biosciences).

### Transmission electron microscopy

bdEV preparations (10 μL) were adsorbed to glow-discharged 400 mesh ultrathin carbon-coated grids (EMS CF400-CU-UL) for 2 minutes following our published protocol (Huang et al., 2020a). Grids were rinsed 3 times with tris-buffered saline and stained with 1% uranyl acetate with 0.05% Tylose. Grids were aspirated, dried, and immediately imaged with a Philips CM120 instrument set at 80 kV. Images were captured with an 8-megapixel AMT XR80 charge-coupled device.

### bdEV and BH RNA extraction

bdEV RNAs were extracted by adding Trizol LS (Thermo Fisher 10296028) to 100 μL bdEV resuspension. After phase separation, RNAs were purified by miRNeasy Mini Kit solutions (Qiagen217004) and Zymo-Spin I Columns (Zymo ResearchC1003–50) according to the manufacturer’s instructions.

BH RNA was extracted by adding Trizol (Thermo Fisher15596018) and homogenizing tissues with Lysing Matrix D (MP Biomedicals116913100) in a benchtop homogenizer (FastPrep-24, MP Biomedicals) at 4.0 m/s for 20 seconds. After homogenization, the supernatant was collected, and RNA was isolated by miRNeasy Mini Kit solutions (Qiagen217004) and Zymo-Spin IIICG Columns (Zymo Research C1006-50-G) following the manufacturer’s instructions.

### SIV Gag RNA quantification by qPCR/ddPCR

Viral RNA was measured by quantitative PCR (qPCR) or digital droplet PCR (ddPCR) after reverse transcription as described (Abreu et al., 2019; Shen et al., 2003). Viral RNA from parietal cortex tissues and CSF was isolated by QIAamp Viral RNA Mini kit (Qiagen 1020953). qPCR of SIV gag RNA was by QuantiTect Virus kit (Qiagen 211011) or ddPCR using One-Step RT ddPCR Advanced Kit for Probes (Bio-Rad 1864022). Copy numbers were calculated with a regression curve from control RNA transcript standards and normalization to per µg RNA in cortex or per ml CSF. Primers/probes for SIV gag RNA were: SIV21 forward 5’-GTCTGCGTCATCTGGTGCATTC-3’; SIV22 reverse 5’-CACTAGGTGTCTCTGCACTATCTGTTTTG-3’; SIV23, 5’ FAM/3’-Black hole quencher-labeled probe 5′-CTTCCTCAGTGTGTTTCACTTTCTCTTCTG-3 (Integrated DNA Technologies).

### Small RNA sequencing

8 µl of bdEV (from a total of 40 μl RNAs) and 20 ng of BH RNAs were used for small RNA library construction by the D-Plex Small RNA-seq Kit (Diagenode C05030001). D-Plex Single Indexes for Illumina - Set A (DiagenodeC05030010) were attached according to the manufacturer’s protocol. The yield and size distribution of the small RNA libraries were assessed using the Fragment Bioanalyzer™ system with DNA 1000 chip (Agilent5067-1505). 170-230 bp libraries were selected with agarose gel cassettes (Sage Science HTG3010) on the BluePippin Size Selection System (Sage Science). Multiplexed libraries were equally pooled to 1nM and sequenced on the NovaSeq 6000 system (Illumina) with the NovaSeq 6000 SP Reagent Kit v1.5 (100 cycles) (Illumina 20028401).

### Small RNA sequencing data analysis

Raw reads were first trimmed from polyA-tails using cutadapt software, and the PCR duplicates were removed by collapsing identical sequences with seqkit. De-duplicated reads were trimmed from the 5’-UMI sequences and size-selected using cutadapt software. Trimmed and size-selected (>15 nt) reads were aligned to custom-curated hg38 reference transcriptomes using Bowtie, allowing 1 mismatch tolerance (-v 1 option) as follows. First, reads were mapped to RNA species with low sequence complexity and/or high repeat number: rRNA, tRNA, RN7S, snRNA, snoRNA, scaRNA, vault RNA, RNY, and mitochondrial chromosome (mtRNA). Unmapped reads were aligned sequentially to mature miRNA, pre-miRNA, protein-coding mRNA transcripts (mRNA), and long non-coding RNAs (lncRNAs). Unmapped reads were aligned to the remaining transcriptome (other ncRNAs: mostly pseudogenes and non-protein-coding parts of mRNAs). Finally, all remaining unmapped reads were aligned to the human genome reference (rest hg38) corresponding to introns and intergenic regions. Data scaling was done using R/Bioconductor packages DESeq2, and scaled data were visualized with principal component analysis (PCA). Differential gene expression was quantified using R/Bioconductor packages DESeq2 (Love et al., 2014) and edgeR (Smyth et al., 2018) with sequences identified by both packages defined as differentially expressed (FDR adjusted p-value < 0.05). RefFinder (Xie et al., 2012) was used for evaluating and screening internal reference genes from sRNA sequencing datasets. mRNA/protein interaction, cluster mRNA/protein function prediction, and cellular component annotations were done by Protein-Protein Interaction Networks Functional Enrichment Analysis (STRING) (Szklarczyk et al., 2019). Hierarchical clustering of sRNAs was performed with Heatmapper (Babicki et al., 2016). miRNA-mRNA interaction predictions were conducted by TargetScan (McGeary et al., 2019) and miRDB (Chen and Wang, 2020). To assess whether SIV infection-associated sRNA differences in the extracellular space reflect overall changes in the brain tissue, Pearson’s correlation was used to evaluate the correlation of sRNA fold changes in bdEVs with BH. Two-tailed p < 0.05 was considered statistically significant. Analysis was conducted in R 4.2.1 and GraphPad Prism.

### Total transcriptome sequencing

1 µg of BH RNAs was incubated with 1 U/µg RNase R (Lucigen RNR07250) at 37°C for 30 min. RNase R-treated RNAs were then re-isolated by RNA Clean & Concentrator™-5 (Zymo Research R1014) according to the manufacturer’s protocol. 100 ng of total BH RNA with and without RNase R treatment was then used to construct cDNA libraries by Illumina Stranded Total RNA Prep Ligation with Ribo-Zero Plus (Illumina 20072063). A Ribo-Zero Plus kit was used to deplete ribosomal RNA in these samples, and library construction was then done per manufacturer’s protocol. Indexes were attached using IDT for Illumina RNA UD Indexes according to the manufacturer’s protocol. The yield and size distribution of the total RNA libraries were assessed using the Fragment Bioanalyzer™ system with DNA 1000 chip (Agilent 5067-1505). Multiplexed libraries were equally pooled to 1nM and sequenced with the NovaSeq 6000 system (Illumina) and NovaSeq 6000 S2 Reagent Kit v1.5 (300 cycles) (Illumina 20028314).

### Total transcriptome sequencing data analysis

The raw reads were first trimmed from contaminating adapter sequences using cutadapt software. The trimmed and size-selected (>15 nt) reads were then aligned using Bowtie2 (default settings) to the manually curated M. mulatta mRNA reference containing a single (main) transcript per each gene (designated with a gene symbol, e.g., GAPDH). All reads which did not align to the above mRNA were mapped to the combined M. mulatta cDNA and ncRNA references from Ensembl (designated with transcript ID, e.g., ENSMMUT00000047080.3). The numbers of reads mapped to each transcript were extracted using eXpress software based on a previous publication (Roberts and Pachter, 2013).

### circRNA identification by Sequential Alignment (CiRISeqA)

All reads aligned to the *M. mulatta* combined cDNA and ncRNA references from Ensembl were discarded. The remaining reads were additionally filtered from non-circular RNA sequences by removing all reads mapped to "non-modified" *M. mulatta* CircAtlas v.2 references (downloaded from: http://circatlas.biols.ac.cn/) using the same Bowtie settings. Next, the remaining reads were aligned to "duplicated" CircAtlas v.2 reference transcripts that correspond to sequences mapped to the junction regions of circRNAs. The numbers of reads mapped to each duplicated RNA reference were extracted using eXpress software based on a previous publication (Roberts and Pachter, 2013). See also Supplementary Figure 2.

### circRNA identification by circExplorer2

The genome and gene annotations for *M. mulatta* were obtained from UCSC (BCM Mmul_8.0.1/rheMac8). First, raw files were aligned to the genome using STAR (chimSegmentMin 10), and the chimeric junction files were used to count circRNAs using circExplorer2. circRNAs identified by both our custom pipeline (above) and circExplorer2 were included in the following analysis.

### circRNA data analysis

Differential circRNA expression was quantified using R/Bioconductor packages edgeR. CircRNAs with false discovery rate (FDR) < 0.05 were defined as significantly differentially abundant. Significant circRNAs from both CiRISeqA and circExplorer2 were included in the analysis. miRanda (Betel et al., 2010) (MMU) was used to predict interactions of circRNAs and mature miRNA sequences (912 macaque miRNAs, miRBase).

### Individual qPCR assays for miRNAs

Individual qPCR assays (Thermo Fisher) were performed as described (Witwer et al., 2011) for miRs-19a-3p (Assay ID 000395), 29a-3p (002112), 146a-5p (000468), 449a-5p (001030), and let-7a-5p (000377). Data were adjusted to the geometric mean of Cq values of selected internal reference genes: miRs-124-3p (003188_mat), 125b-5p (000449), and 23b-3p (245306_mat).

### Individual qPCR assays for circular RNAs

1 µg BH RNAs and 5 µl bdEV RNAs were used to generate cDNA by iScript ™ Reverse Transcription Supermix for RT-qPCR (Bio-Rad 1708840). Expression levels of circRNAs were evaluated by qPCR using iTaq Universal SYBR ® Green Supermix (Bio-Rad 1725120). Divergent primers for circRNAs were designed using the primer3 webtool (Untergasser et al., 2012). The 3’ end of a circRNA is fused to its 5’ end and then submitted to primer3 for the primer design using default parameters. GAPDH (forward 5’-CCATGGGGAAGGTGAAGGTC-3’, reverse 5’-TGAAGGGGTCATTGATGGCA-3’) was used as a housekeeping gene for circRNA quantification in BH, while the geometric mean of Cq values of selected internal reference genes (miRs-124-3p, 125b-5p, and 23b-3p) was used as a reference for circRNA in bdEVs.

### Statistical analysis, data availability, and EV-TRACK

Statistical significance of particle count, size distribution differences, miRNA level, and circRNA level between different groups were determined by the Brown-Forsythe and Welch ANOVA tests.

We have submitted all relevant details of our experiments to the EV-TRACK knowledgebase (EV-TRACK ID: EV230365). Nucleic acid sequencing data (currently being deposited with NCBI databases and available on request during review).

## Results

Following the protocol illustrated in Figure 1A, bdEVs were separated from the occipital brain tissue of uninfected and SIV-infected macaques. After basic bdEV characterization, bdEVs and source brain homogenate (BH) were subjected to small RNA-Seq for mRNA and miRNA identification, while BH was subjected to parallel total RNA-Seq for mRNA and circRNA identification.

**Figure 1.**
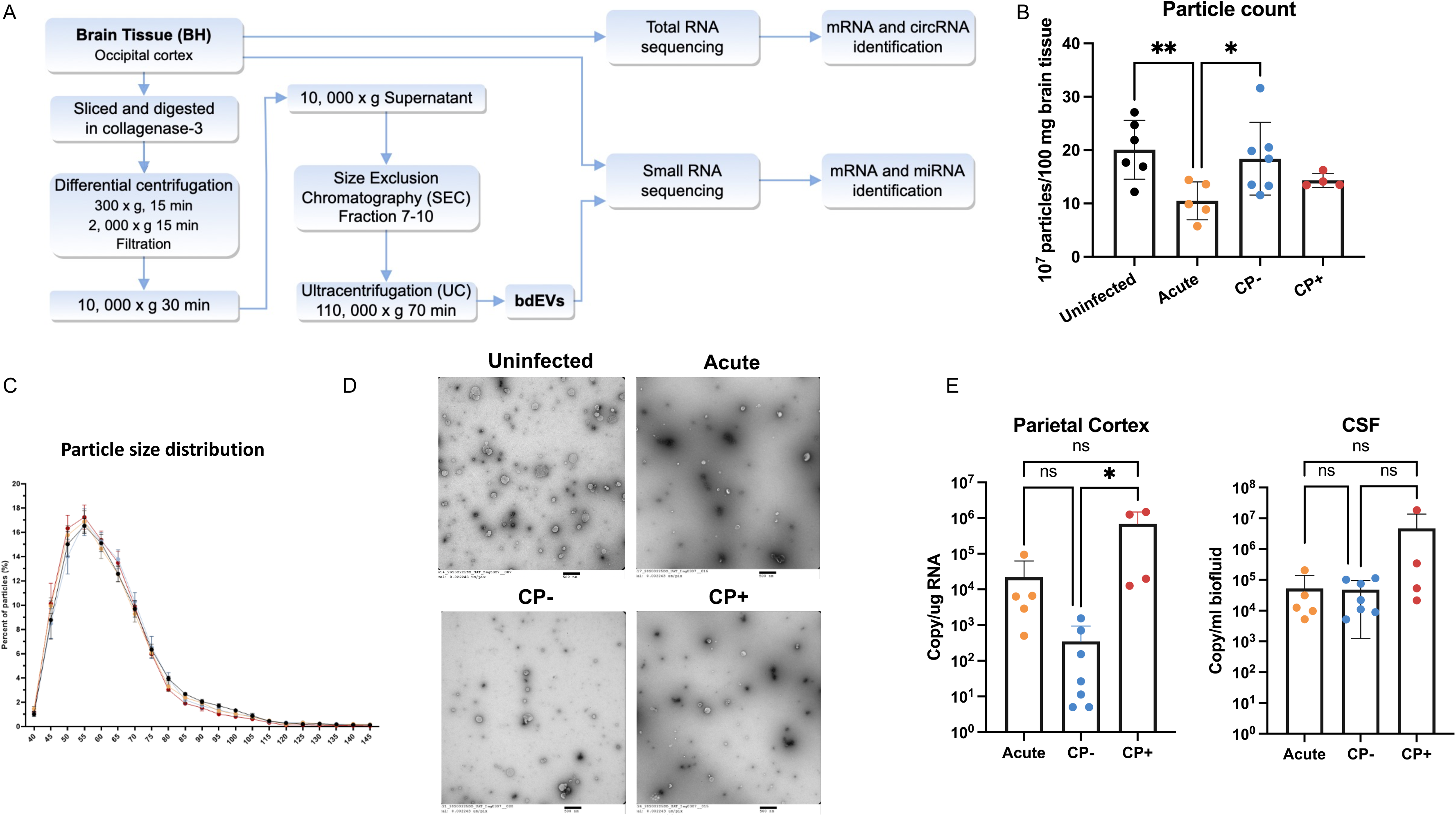
Enrichment and characterization of brain tissue-derived EV (bdEV) from uninfected and SIV-infected macaques. A) Workflow for bdEV enrichment and RNA sequencing. bdEV and source brain homogenate (BH) were subjected to small RNA-Seq, while BH was subjected to parallel total RNA-Seq. B) Particle concentrations of bdEVs from uninfected, acute, CP-, and CP+ macaques were measured by NFCM. Particle concentration for each group was normalized by tissue mass (per 100 mg). Data are presented as mean ± SD. *p≤0.05, **p≤0.01 by two-tailed Welch’s t-test. C) Size distributions of bdEVs were measured by NFCM and calculated as particles in a 5nm size bin versus total detected particles in each sample (percentage). D) bdEVs were visualized by negative staining transmission electron microscopy (TEM) (scale bar_=_500 nm). TEM is representative of ten images taken of each fraction from three independent human tissue samples. E) Viral GAG RNA amplicon as copy number per ug RNA in parietal cortex, and per ml CSF measured by qPCR. Data are presented as mean ± SD. ns, not significant, *p≤0.05 by two-tailed Welch’s t-test.

### Particle counts, sizes, and morphology in SIV infection

Particle count and size distribution per 100_ mg tissue input were determined for bdEV preparations by NFCM. Fewer particles were recovered from acute-phase samples compared with the uninfected and chronic infection without CNS pathology (CP-; Figure 1B). No overall particle size distribution difference was detected by NFCM (Figure 1C). Transmission electron microscopy (TEM) revealed oval and round particles, consistent with EV morphology (Figure 1D). EV-associated membrane proteins CD81, CD63, and CD9 were detected by single-particle interferometric reflectance imaging sensing (SP-IRIS) (Supplementary Figure 1). To assess viral replication, an SIV RNA gag amplicon was measured by qPCR in parietal cortex tissue and CSF corresponding to each brain sample (Figure 1E). More gag amplicons were observed in the CP+ group compared with the CP-group in parietal cortex, while no statistically significant differences between groups were observed in CSF.

### bdEV sRNA dysregulation in SIV infection and CP+

Ligation-independent small RNA-Seq of bdEVs yielded an average of 38.7M (± 5.4M) reads per sample (M = million, 1 x 10^6). After adapter clipping and removing reads shorter than 15 nt, 85.04% (± 1.81%) of bdEV reads mapped to the human genome (hg38). PCA based on sRNA profiles (Figure 2A) indicated a clear separation of the acute group. In the CP+ group (n=4), sRNA contents of three bdEV samples were separated from the other groups, while one was close to the CP-group. Differential expression analysis was conducted to identify differentially abundant sRNAs in SIV versus uninfected and CP+ versus CP-based on adjusted p-value < 0.05 (Figure 2B). Most differences were between acute and uninfected (71 more, 6 less abundant), followed by CP+ versus uninfected (46 more, 2 less abundant), and CP+ versus CP-(2 more abundant). No significantly different sRNAs were identified in CP-versus uninfected. Among all dysregulated sRNAs, 38 were consistently dysregulated in both acute and CP+ versus uninfected, while 2 were also dysregulated in CP+ versus CP-(Figure 2C). As all 38 sRNAs were mapped to protein-coding messenger RNAs (mRNAs), STRING analysis was used to examine the potential functions of their parent mRNAs. 30 out of 38 mRNAs had a high interaction confidence score as predicted by STRING (0.7 on a scale of 0-1) (Figure 2D). Gene ontology (GO) enrichment analysis indicated potential involvement in immune regulation, including cytokine-mediated signal pathways, interferon-gamma mediated signal pathways, immune system process, immune effector process, and defense responses to viruses. (Figure 2E). To examine the associations of CP+ and bdEV sRNA levels, 10 sRNAs with putative differential abundance exclusively between CP+ and uninfected (Figure 2D, sRNAs indicated in the upper right of the Venn diagram) were then visualized by unsupervised clustering (Figure 2F). Based on the profile of these sRNAs, subjects in CP+ and uninfected groups clustered together (Figure 2F). Although no significant GO ontology enrichment was found for these sRNAs (data not shown), many have known involvement in immunity and immune regulation based on the Database for Annotation, Visualization and Integrated Discovery (DAVID) (Sherman et al., 2022), including protein tyrosine phosphatase (PTPRC), complement C7 (C7), IFI6 interferon alpha inducible protein 6 (IFI6), CD74, and HLA-DRA.

**Figure 2.**
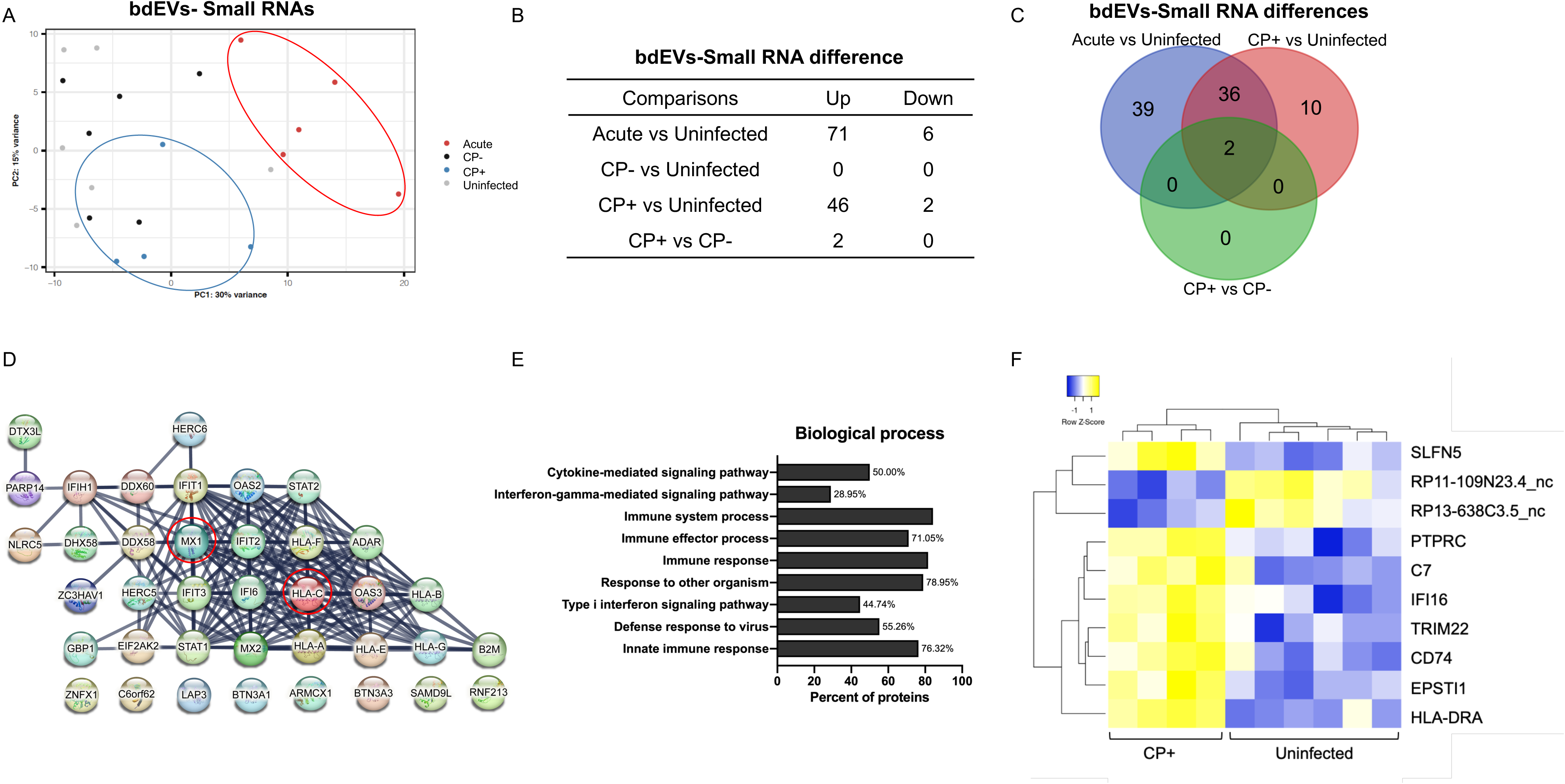
bdEV small RNAs with differential expression in SIV infected macaques. A) Principal component analysis (PCA) based on quantitative small RNA profiles of bdEVs from uninfected, acute, CP-, and CP+ groups. Animal #299 in the CP-group was defined as outlier and not shown in the PCA plot as it was not well reconstructed within the principal vectors. B) The number of small RNAs differentially abundant between groups based on adjusted p-value < 0.05 from both DESeq2 and edgeR packages. C) Venn diagrams of differentially abundant small RNAs in acute and CP+ as compared with uninfected, and CP+ as compared with CP-. D) STRING protein interaction network analysis indicated that 30 out of 38 mRNAs dysregulated in both acute and CP+ groups had a high interaction confidence score (0.7 on a scale of 0-1). E) Top 10 biological processes ranked by FDR-corrected p-value was predicted by GO ontology enrichment for 38 mRNAs dysregulated in both acute and CP+ groups. F) Unsupervised hierarchical clustering of 10 differentially abundant small RNAs of bdEVs between CP+ and uninfected.

### bdEV miRNA dysregulation in SIV infection and CP+

Since we previously reported miRNA dysregulation in biofluids during retroviral infection (Huang et al., 2023; Witwer et al., 2011; Zhao et al., 2020), and since our sample size was relatively small, we further assessed differentially abundant bdEV miRNAs based on unadjusted p-value alone. In line with differences for other sRNAs, more miRNA differences were apparent in the comparison of acute versus uninfected (n=24) than in the CP+ versus uninfected (n=14), or CP+ versus CP-groups (n=9) (Figure 3A and B). Only 3 miRNAs were dysregulated in both acute and CP+ versus uninfected, while 3 miRNAs were dysregulated in both CP+ versus uninfected and CP+ versus chronic groups (Figure 3B). Unsupervised clustering of 20 CP+-associated miRNAs suggested two miRNA clusters that differentiate CP+ and uninfected (Figure 3C). miRNA profiles of CP-are intermediate, with greater variance between subjects. Individual qPCR assays confirmed that miR-19a-3p, let-7a-5p, and miR-29a-3p were less abundant during SIV infection (acute group), and that miRs-146a-5p and -449a-5p were dysregulated in the CP+ group (Figure 3D).

**Figure 3.**
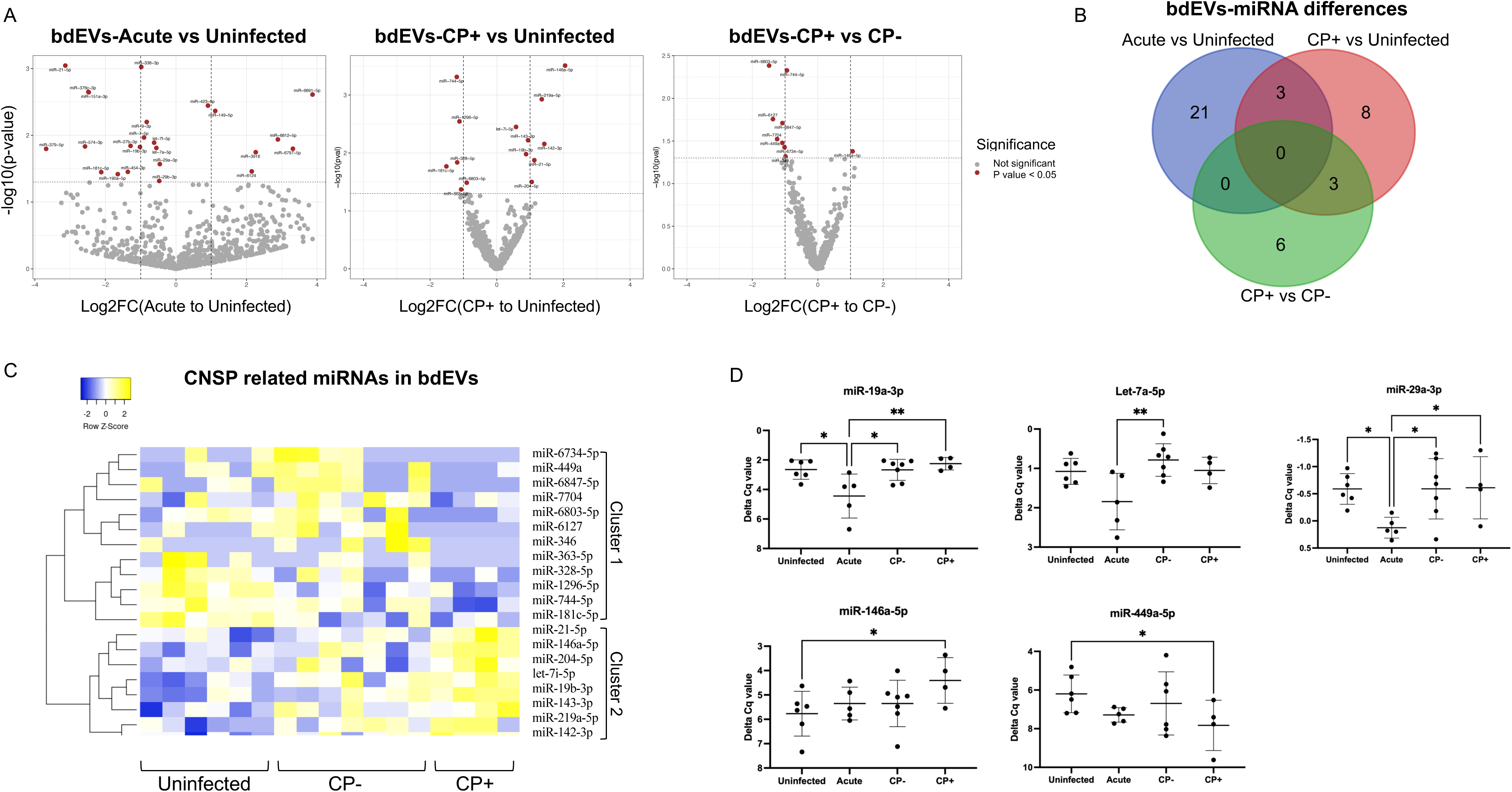
bdEV miRNAs with differential expression in SIV infected macaques. A) Volcano plots showing bdEV miRNA log2FC and p-value for acute vs uninfected (left), CP+ vs uninfected (middle), and CP+ vs CP-(right). Thresholds for 2-fold change and p-value < 0.05 are indicated by dashed lines. Significant changes are indicated with different colors. Gray: Not Significant, Red: non adjust p-value < 0.05. B) Venn diagrams of differentially abundant miRNAs (non-adjust p-value < 0.05) in acute and CP+ as compared with uninfected, and CP+ as compared with CP-. C) Unsupervised hierarchical clustering of 20 miRNAs differentially abundant in bdEVs of CP+ as compared to CP- and uninfected. D) qPCR validation for miR-19a-3p, let-7a-5p, miR-29a-3p, miR-146a-5p, and miR-449a-5p in bdEVs from uninfected, acute, CP-, and CP+ groups. Delta Cq values was normalized to the geometric mean Cq value of selected internal references: miR-124-3p, miR-125b-5p, and miR-23b-3p. Data are presented as mean +/- SD. *p ≤ 0.05, **p ≤ 0.01 by two-tailed Welch’s t-test.

### circular RNA dysregulation in SIV-infected brain

After ribosomal RNA depletion, RNAs from bdEVs and source BH were used to make test libraries for total RNA-Seq. High-quality libraries with expected insert sizes were obtained only from BH-derived RNA, consistent with previous observations that RNAs in EVs are mostly shorter fragments (data not shown). BH RNA was then used for total RNA sequencing. RNase R-untreated and RNase R-treated RNAs were sequenced to identify mRNAs and circRNAs.

PCA was conducted on mRNA and circRNA profiles of BH from uninfected, acute-phase, CP-, and CP+ groups (Figure 4A). Similar to the bdEV sRNA patterns, both mRNA and circRNA profiles in the acute BH group showed separation from other groups. mRNA and circRNA profiles in the uninfected group also separated from the three infected groups, indicate an influence of SIV infection on RNA profiles (Figure 4A). According to the differential expression analysis, in line with bdEVs, most differentially expressed mRNAs in BH were found in the comparison of the acute and uninfected (n=245), followed by CP+ versus uninfected (n=78) (Figure 4B, left). No significantly different mRNAs were identified in either CP+ versus CP- or CP-versus uninfected. 69 mRNAs were consistently dysregulated in both acute and CP+ versus uninfected (Figure 4B, left). For circRNAs, most differentially abundant circRNAs in BH were also found in the comparison of the acute and uninfected (n=19), followed by CP+ versus uninfected (n=10), CP-versus uninfected groups (n=10), and CP+ versus CP-(n=4) (Figure 4B, right). However, only a few circRNAs were identified in more than one comparison (Figure 4B, right). Furthermore, the linear RNAs corresponding to most of the differentially expressed circRNAs did not change significantly in SIV infection or CP+ (Figure 4C, grey dots). An exception was the linear transcript IFI6, which was positively correlated with circ-IFI6_0001 (Figure 4C, shown in red dots). Individual qPCR assays confirmed dysregulation of circ-IFI6_0001, circ-EXOC2_0008, circ-PRKCE_0004, circ-PPP2R5A_0001, circ-RNF41_0003, and circ-ENC1_0001 in SIV-infected BH (Figure 4D).

**Figure 4.**
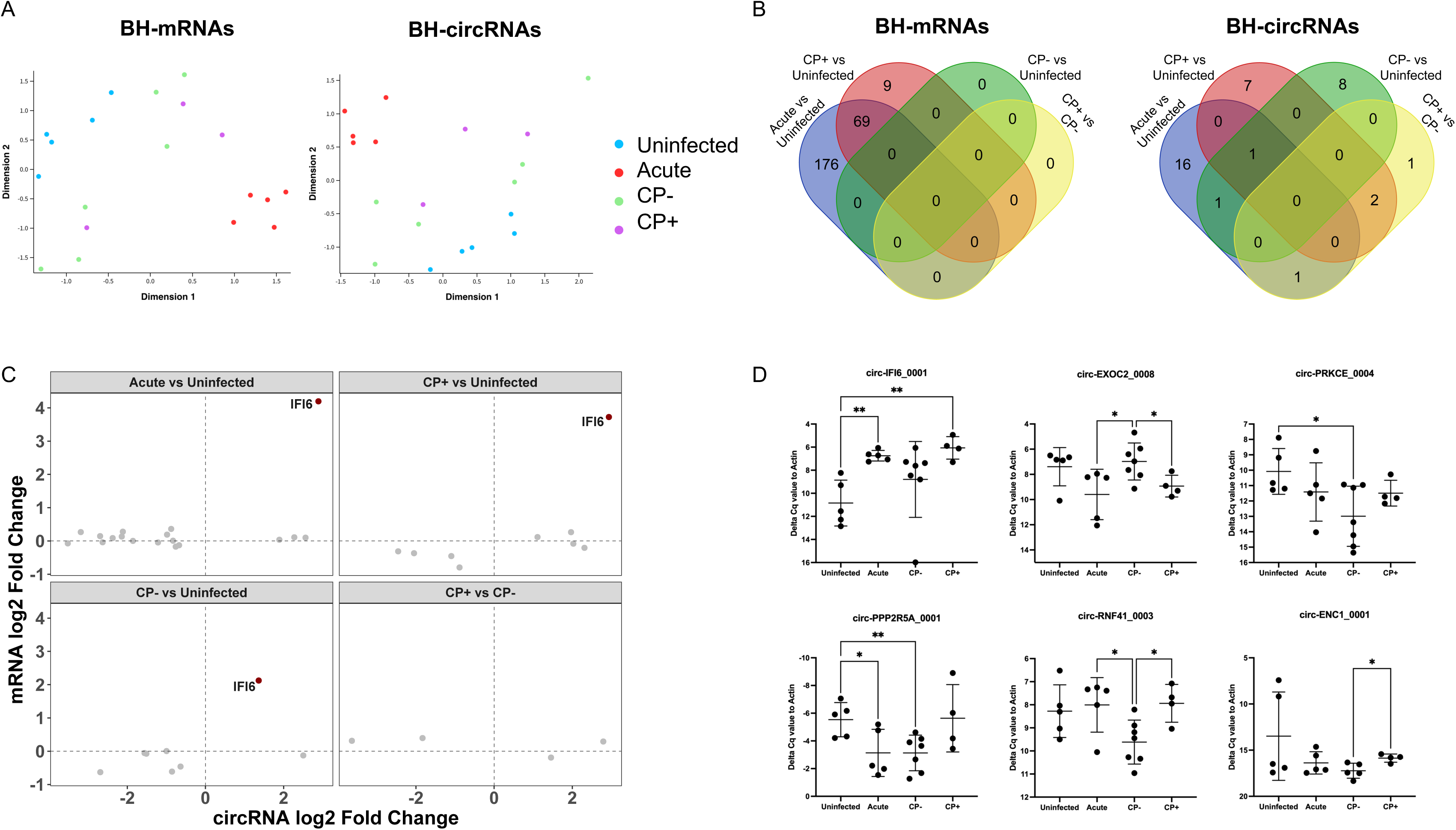
circRNA dysregulation in SIV-infected brain tissues. A) Multidimensional scaling analysis based on quantitative mRNA (left) and circRNA (right) profiles of BH from uninfected, acute, CP-, and CP+ groups. B) Venn diagrams of differentially abundant mRNAs (left) and circRNAs (right) (adjust p-value < 0.05) in acute, CP+, and CP-groups as compared with uninfected, and CP+ as compared with CP-. C) circRNA log2FC between different comparisons in BH were plotted against the correspondent linear mRNA log2FC. Red dots indicate both circRNA and corresponding linear RNA were differentially abundant between groups with adjust p-values < 0.05. D) qPCR validation for circ-IFI6_0001, circ-EXOC2_0008, circ-PRKCE_0004, circ-PPP2R5A_0001, circ-RNF41_0003, and circ-ENC1_0001 in BH from uninfected, acute, CP-, and CP+ groups. Delta Cq values was normalized to the Cq value of GAPDH. Data are presented as mean +/- SD. *p ≤ 0.05, **p ≤ 0.01 by two-tailed Welch’s t-test.

### bdEVs reflected mRNA and circRNA changes in BH after SIV infection

To assess whether SIV infection-associated sRNA differences in the extracellular space reflect overall changes in the brain tissue, we assessed the correlation of sRNA fold changes in bdEVs with BH by Pearson’s correlation analysis (Figure 5A and 5B). sRNAs significantly different (adjusted p-value < 0.05) in both bdEVs and BH when comparing acute with uninfected (n=43, Figure 5A) and CP+ with uninfected (n= 22, Figure 5B) were included in the analysis. A significantly positive correlation of sRNA profiles between bdEVs and BH was observed for both acute vs uninfected (R=0.836, p<0.001) and CP+ vs uninfected (R=0.647, p=0.001) (Figures 5A and 5B). When comparing sRNA profiles between CP+ and CP-, two differentially abundant sRNAs, HLA-C and MX1, were identified only in bdEVs but not in BH. However, the pattern was consistent between bdEVs and BH, with a higher abundance in acute and CP+, and lower in uninfected and CP-(Supplementary Figure 3).

**Figure 5.**
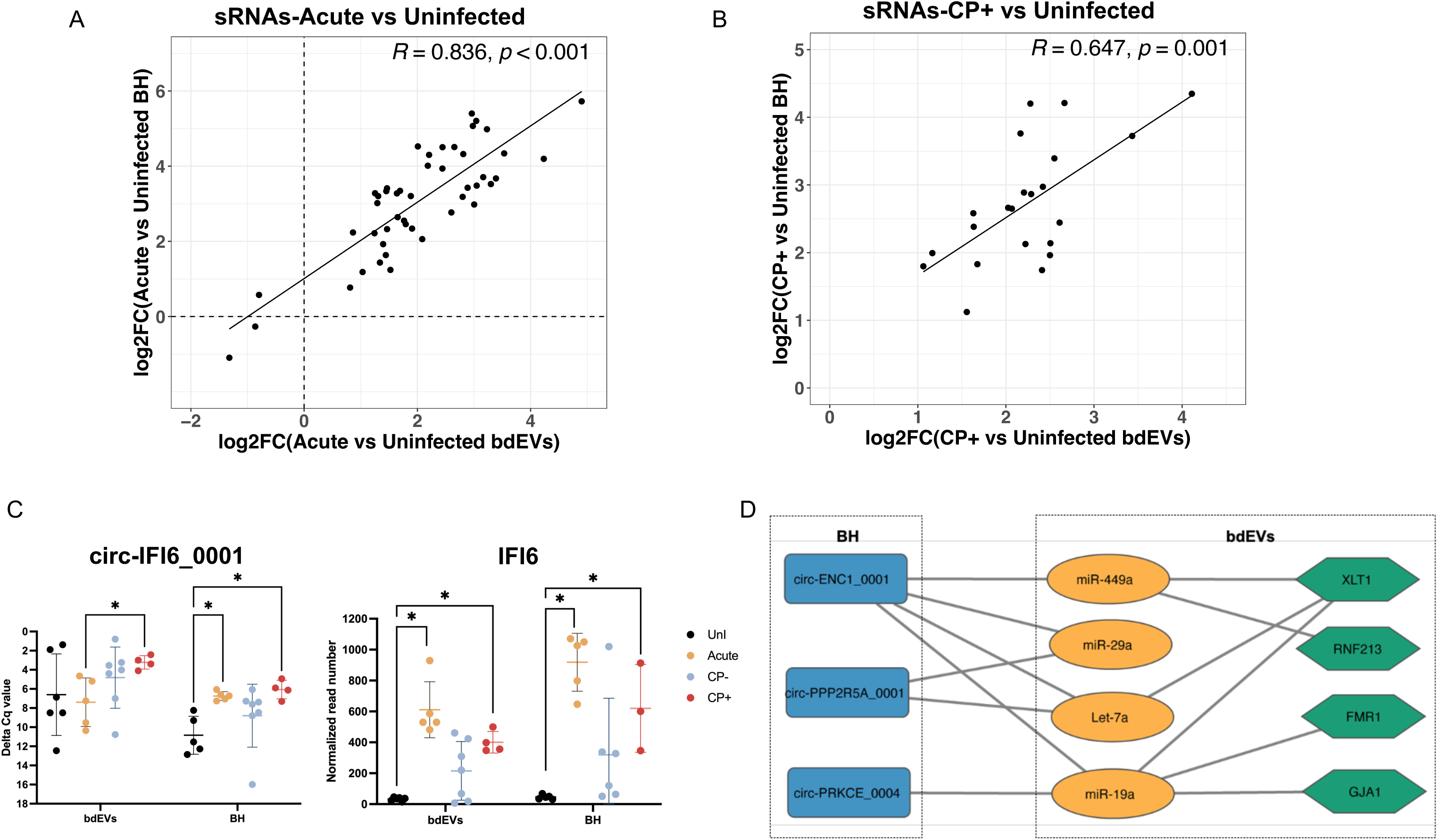
The SIV related mRNA and circRNA changes in bdEVs and BH. A) Correlations of small RNA log2FC in bdEVs and BH when comparing acute with uninfected. Linear regression line was shown in black. Pearson correlation coefficient (R) and significance (p) are shown based on all dysregulated small RNAs in acute versus uninfected. B) Correlations of small RNA log2FC in bdEVs and BH when comparing CP+ with uninfected. Linear regression line was shown in black. Pearson correlation coefficient (R) and significance (p) are shown based on all dysregulated small RNAs in CP+ versus uninfected. C) Levels of circ-IFI6_0001(left) and IFI6 (right) in bdEVs and BH. *adjust p < 0.05 based on edge R. D) The circRNA-associated ceRNA network. The blue rectangle nodes represent circRNAs, the orange oval nodes represent miRNAs, the green hexagon nodes denote mRNAs. The edges represented interactions between RNAs in the network.

circRNA differences in bdEVs were also tested by individual qPCR assays. Among six dysregulated circRNAs identified in SIV-infected BH (as shown in Figure 4D), only circ-IFI6_0001 was consistently more abundant in CP+ for both bdEVs and BH (Figure 5C, left). The corresponding linear transcript, IFI6, was also more abundant in acute and CP+ compared with uninfected (sequencing data), and for both bdEVs and BH (Figure 5C, right).

To identify potential ceRNA regulation networks involving bdEVs and BH, bdEV miRNAs (Figure 3D) and BH circRNAs (Figure 4D) verified by qPCR were used as input for target prediction, along with mRNAs. circRNAs, miRNAs, and mRNAs were represented as nodes, while the edges represented interactions between items in the network (Figure 5D). miRanda predicted that three circRNAs in BH (Figure 5D, blue rectangle) potentially bind with four miRNAs identified in bdEVs (Figure 5D, orange ellipse). Next, overlapping mRNA targets of these four miRNAs were predicted by TargetScan and miRDB (Supplementary Figure 4). Based on the overlapping mRNA list and differences in bdEVs, four potential mRNA targets are indicated in Figure 5D (green hexagon).

## Discussion

The potential regulatory roles of ceRNA networks in HIV/SIV-infected brains and neuroinflammation are largely undetermined, for both brain tissue and extracellular particles like bdEVs that may mediate communication between cells. Here, we systematically examined RNA profiles of bdEVs and source brain tissues of SIV-infected HIV models. Overall bdEV sRNA profile and brain tissue transcriptome differences were most pronounced for the acute infection group, suggesting that retroviral infection has the greatest effect on host RNA processing in brain early in infection. However, some differences were also observed in association with CNS pathology in chronic infection. GO enrichment analysis suggested that dysregulated mRNAs (or fragments thereof) in bdEVs are significantly involved in inflammation regulation and immune responses. Furthermore, competing RNA network analysis showed that several circRNAs in brain cells may affect the miRNA and mRNA contents of bdEVs. ceRNA networks in the brain and bdEVs may contribute to gene expression modulation in HIV/SIV infection and neuropathology. Further studies are needed to confirm and characterize the extent of these associations.

One aspect of our findings regarding the correlation of EV abundance and HIV infection contrasts starkly with earlier reports. Whereas previous studies (Bazié et al., 2021; Chettimada et al., 2020; Hubert et al., 2015; WW et al., 2021), including our own recent study on SIV infection (Huang et al., 2023), report more abundant plasma EVs during SIV infection, especially during acute infection phase, we report lower particle concentrations in bdEV preparations from acute-infected subjects. Virions themselves are unlikely to explain either the EV abundance in plasma or in brain: although virions co-isolate with EVs (Hoen et al., 2016), they are not abundant enough even in acute infection to contribute to overall particle increases in plasma. Instead, the cellular origins and uptake patterns of EVs in the peripheral blood (Auber and Svenningsen, 2022; Grenier-Pleau et al., 2020; Li et al., 2020) and brain compartments (Arab et al., 2022; Huang et al., 2022a) may be responsible for the seemingly discrepant findings. Studies of EV release/uptake dynamics of different cell types and during retroviral infection and inflammation are merited.

Previous studies of cell models revealed host gene expression changes associated with HIV infection and HIV-associated neurocognitive disorders (Chang et al., 2011; Devadas et al., 2016; Ojeda-Juárez and Kaul, 2021; Sanna et al., 2021). Here, we for the first time compared mRNA changes in bdEVs and brain tissues in the SIV/HIV infection model, including differences associated with CNS pathology. We found that acute infection and chronic pathology (CP+) groups shared many common mRNA differences when compared with the uninfected group, while in the CP-group, no significant differences were detected compared with uninfected group. 8 signature mRNAs in bdEVs were dysregulated in the CP+ group, possibly representing genes that are especially affected in SIV-associated CNS pathology. These genes encode key immune system proteins such as CD74 and HLA-DRA, involved in antigen processing and presentation (Karakikes et al., 2012; Su et al., 2017). Another mRNA was for the interferon γ-inducible protein 16 (IFI16) (Altfeld and Gale, 2015a), which modulates HIV transcription, latency reactivation, and HIV replication by targeting the transcription factor Sp1 to drive viral gene expression (Hotter et al., 2019) or as the immune sensor of retroviral DNA (Altfeld and Gale, 2015b; Jakobsen et al., 2013). These mRNAs are differentially abundant not just in BH, but also in bdEVs (Figure 5A-B), suggesting that CNS cells may use bdEVs to transfer antiviral immune response elements during SIV infection.

In addition to mRNAs, we examined miRNAs, a class of RNA also reportedly altered in HIV infection (Klase et al., 2012; Swaminathan et al., 2014, 2012a). Here, only subtle differences were detectable, but several could still be verified by qPCR. Two miRNAs (let-7a-5p and miR-29a-3a) were previously reported in HIV infection or inflammation. Consistent with a previous report that let-7 family miRNAs were less abundant in CD4+ T cells from HIV-1-infected patients compared with uninfected controls and long-term non-progressors, we found less let-7a-5p in bdEVs from acute infection (Swaminathan et al., 2012b). Since let-7 might regulate host immune responses by increasing IL-10 levels (Swaminathan et al., 2012b) or affecting other chemokine and cytokine-related pathways (Venkatachari et al., 2017) in peripheral blood cells, it might also function in SIV CNS infection by similar pathways. miR-29a, which we previously also found to be less abundant in peripheral plasma EVs during acute infection (Huang et al., 2023) may directly target many regions of HIV RNA to inhibit HIV viral translation and replication (Ahluwalia et al., 2008; Nathans et al., 2009; Sun et al., 2012b), even during HIV latency (Patel et al., 2014a, 2014b). We also identified two miRNAs correlated with SIV-associated CNS pathology, including neuroinflammation-related miRNA-146a-5p (Kim et al., 2022; Rom et al., 2010; Zhao et al., 2023) and brain development-related miR-449a-5p(Tan et al., 2020; Wu et al., 2014). Furthermore, let-7a, miR-29a, and miR-449a were key components of a ceRNA network we constructed between BH and bdEVs (Figure 5D), indicating that their levels in bdEVs, as well as those of target mRNAs, may be regulated by upstream circRNA expression in brain tissue.

We also profiled circRNAs in SIV-infected brain tissues. Specific circRNAs have differential abundance not only in the acute and CP+ groups, but also in the CP-group compared with controls. These circRNAs include circ-PRKCE_0004, circ-PPP2R5A_0001, and circ-RNF41_0003, as also verified by qPCR assays, indicating persistent RNA dysregulation even in asymptomatic chronic infection. We also found two circRNAs, circ-IFI6_0001 and circ-EXOC2_0008, with levels consistent with dynamic changes during the course of SIV infection and disease. The level of circ-IFI6_0001 was greater in the acute and CP+ groups but had a similar level in uninfected and CP-groups, while circ-EXOC2_0008 showed the reverse pattern. These differences indicate potential regulatory roles and biomarker potential in SIV CNS infection. Among qPCR-verified circRNAs, circ-PRKCE_0004 and circ-ENC1_0001 were part of the ceRNA networks we identified that may potentially affect the level of several miRNAs in bdEVs. Regulatory roles of these covalently closed forms of RNAs in viral infection and CNS diseases are largely unexplored, but functions of their linear forms are known. IFI6 is an interferon-induced proteins with antiviral effects against HIV, influenza, and flavivirus (Jiang et al., 2021; Kubo et al., 2022; Richardson et al., 2018). Exocyst Complex Component 2 (EXOC2), involved in vesicle-mediated transport, has critical roles in vesicle tethering and fusion with the plasma membrane but also in neuronal function and brain development (Van Bergen et al., 2020). Consistent differential abundance of circular and linear IFI6 in both SIV-infected BH as well as in bdEVs emphasizes the potential roles of bdEVs in transferring viral infection regulators in brain. Overall, although several groups have reported that different viral infections affect circRNA expression patterns [e.g., (Xie et al., 2021)], only one publication to our knowledge has investigated circRNA in HIV infection (Zhang et al., 2018b). Our findings thus provide additional evidence of circRNA dysregulation in retroviral infection. Further studies are now needed to determine the regulatory mechanisms of these circRNAs in SIV CNS infection and assess their biomarker or therapeutic potential.

In conclusion, RNA profiling of bdEVs and source brain tissues from the SIV model of HIV CNS disease identified mRNAs, miRNAs, and circRNAs closely linked to SIV infection and neuropathology in both bdEVs and BH. Our study provides additional evidence of ceRNA network dysfunction in SIV-related CNS disease. Our findings also have several limitations. First, the size of the study was relatively small. Second, the SIV model may not entirely recapitulate all aspects of HIV disease in people living with HIV (PLWH). Lastly, only selected number of targets were verified by qPCR assays. Our results should thus be further explored and verified using larger cohorts and in HIV infection to assess theragnostic potential.

---

Supplementary Figure 1 Levels of EVmarkers CD81, CD63, and CD9 were measured by single-particle interferometric reflectance imaging (SP-IRIS) and normalized per 100 mg tissue input.

Supplementary Figure 2 Workflow for circRNA identification by Sequential Alignment (CiRISeqA) pipeline.

Supplementary Figure 3 Levels of mRNA HLA-C(left) and MX1 (right) in bdEVs and BH. *adjust p < 0.05 based on edgeR.

Supplementary Figure 4 mRNAs targets of miRNAs as predicted by TargetScan and miRDB. Venn diagrams of mRNA targets as predicted by TargetScan and miRDB for miR-449a, miR-29a-ap, Let-7a-5p, and miR-19a-3p.

## Acknowledgments

The authors thank members of the Witwer Laboratory for discussions and support. We are particularly grateful to members of the Retrovirus Laboratory for access to samples from the animal models and for helpful suggestions. Electron microscopy images were acquired in the Johns Hopkins University School of Medicine Institute for Basic Biomedical Sciences Microscope Facility. RNA sequencing was performed in the Johns Hopkins University School of Medicine Single Cell & Transcriptomics Core. This work was supported in part by the US National Institutes of Health, National Institute on Drug Abuse (NIDA, DA040385 and DA047807 to KWW), by two pilot grants awarded to YH through the Johns Hopkins NIMH Center (supported by MH075673) and the Johns Hopkins University Center for AIDS Research (supported by AI094189), and by NSF 2244127 to MB. The Witwer lab is also supported in part by NCI/Common Fund CA241694, NIAID AI144997, NIMH MH118164, and the Richman Family Precision Medicine Center of Excellence in Alzheimer’s Disease. Samples used in this study were derived in part from research supported by U42OD013117 (Johns Hopkins, NIH supported pigtailed macaque breeding colony), NINDS NS089482 (to Joseph L. Mankowski), and NIMH MH070306 (to Janice E. Clements).

## Author contributions

K.W.W, Y.H. and L.Z. conceived the idea. Y.H. performed most experiments and drafted and revised the manuscript; K.W.W. directed the project, obtained funding, supervised the experiments, and revised the manuscript; A.A, A.T. and M.B. performed small RNA and total RNA sequencing data analysis; S.Q. and C.M.A processed and provided the macaque brain samples and did GAG RAN qPCR; A.T. and X.Z. optimized figure quality in the manuscript. All authors read and approved the final manuscript.

