## Supplemental Figures for "RNA landscapes of brain tissue and brain tissue-derived extracellular vesicles in simian immunodeficiency virus (SIV) infection and SIV-related central nervous system pathology"

Supplementary Figure 1

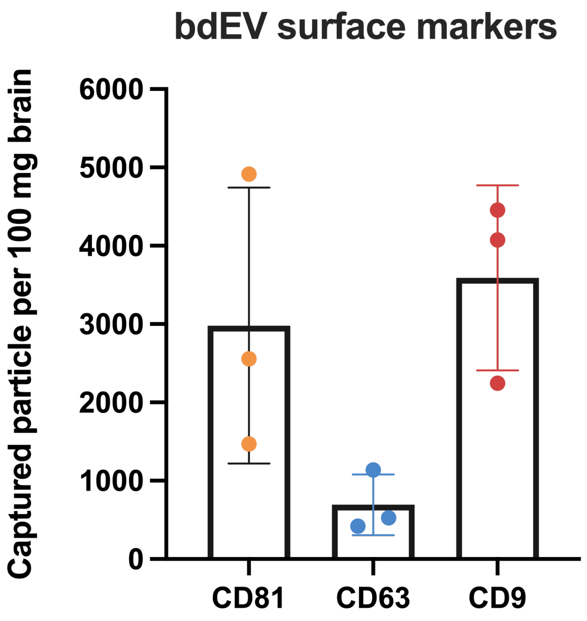

Supplementary Figure 2

#### circRNA identification by Sequential Alignment (CiRISeqA)

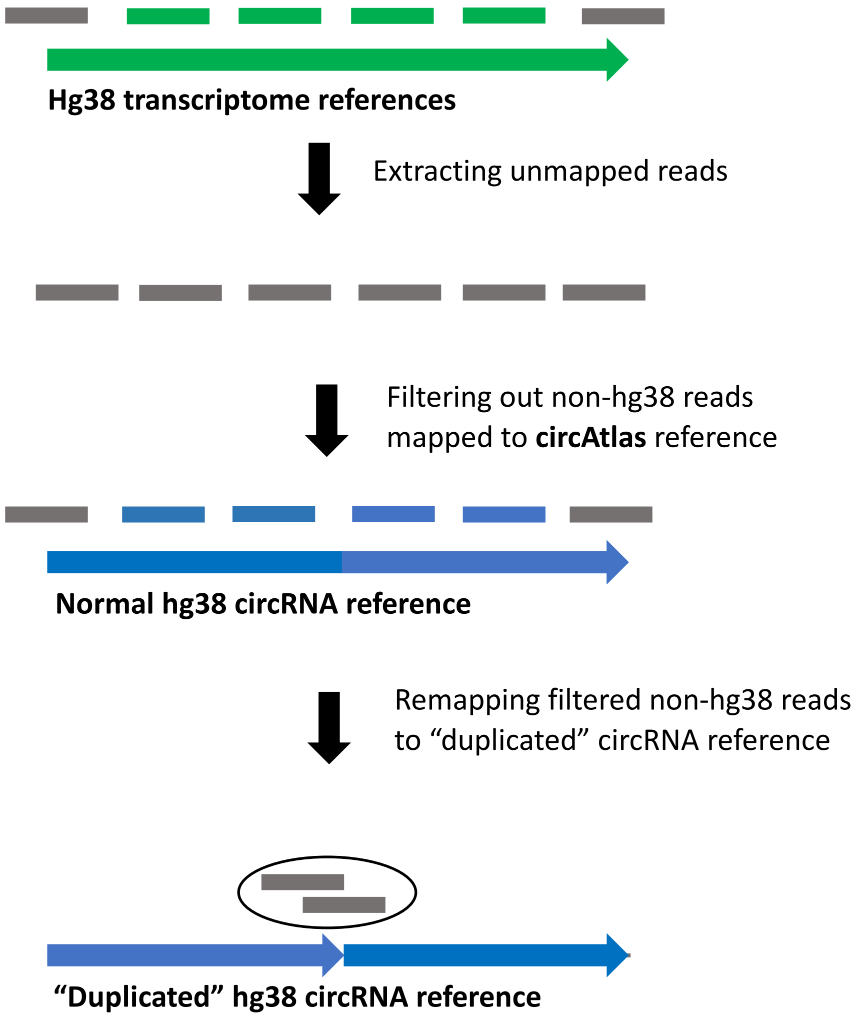


| CircRNA | Read 1 counts |
|---|---|
| hsa-circR-1 | 1000 |
| hsa-circR-2 | 10 |
| hsa-circR-3 | 0 |
| hsa-circR-4 | 5000 |
| hsa-circR-5 | 0 |
| hsa-circR-6 | 200 |

| CircRNA | Read 2 counts |
|---|---|
| hsa-circR-1 | 500 |
| hsa-circR-2 | 0 |
| hsa-circR-3 | 1000 |
| hsa-circR-4 | 5000 |
| hsa-circR-5 | 0 |
| hsa-circR-6 | 100 |

| CircRNA | Read 1+2 counts |
|---|---|
| hsa-circR-1 | 1500 |
| hsa-circR-2 | 10 |
| hsa-circR-3 | 1000 |
| hsa-circR-4 | 10000 |
| hsa-circR-5 | 0 |
| hsa-circR-6 | 300 |

### Supplementary Figure 3

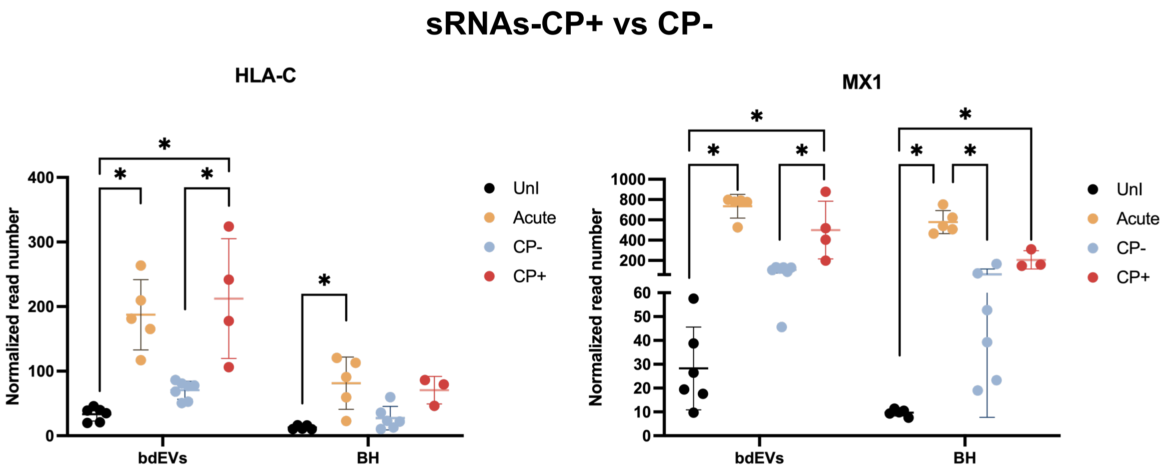

### Supplementary Figure 4

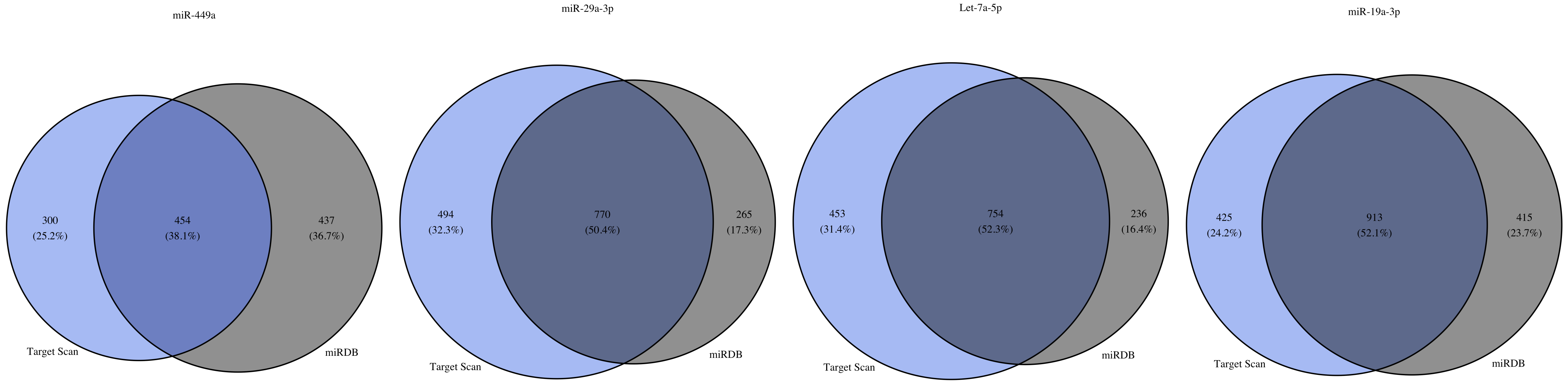
